# Designing Peptides on a Quantum Computer

**DOI:** 10.1101/752485

**Authors:** Vikram Khipple Mulligan, Hans Melo, Haley Irene Merritt, Stewart Slocum, Brian D. Weitzner, Andrew M. Watkins, P. Douglas Renfrew, Craig Pelissier, Paramjit S. Arora, Richard Bonneau

## Abstract

Although a wide variety of quantum computers are currently being developed, actual computational results have been largely restricted to contrived, artificial tasks. Finding ways to apply quantum computers to useful, real-world computational tasks remains an active research area. Here we describe our mapping of the protein design problem to the D-Wave quantum annealer. We present a system whereby Rosetta, a state-of-the-art protein design software suite, interfaces with the D-Wave quantum processing unit to find amino acid side chain identities and conformations to stabilize a fixed protein backbone. Our approach, which we call the *QPacker*, uses a large side-chain rotamer library and the full Rosetta energy function, and in no way reduces the design task to a simpler format. We demonstrate that quantum annealer-based design can be applied to complex real-world design tasks, producing designed molecules comparable to those produced by widely adopted classical design approaches. We also show through large-scale classical folding simulations that the results produced on the quantum annealer can inform wet-lab experiments. For design tasks that scale exponentially on classical computers, the *QPacker* achieves nearly constant runtime performance over the range of problem sizes that could be tested. We anticipate better than classical performance scaling as quantum computers mature.

## Introduction

Computational protein design involves astronomically large search problems. Given *N* designable sequence positions and *D* discrete side-chain identity and conformation possibilities (termed *rotamers*) at each position, there are *D*^*N*^ possible solutions to the problem of finding the optimal selection of one rotamer per position.^1^ Since typical design tasks might involve tens or hundreds of positions, with hundreds or even thousands of possibilities for the rotamer at each position, the naïve, exhaustive approach to these tasks rapidly exceeds the capabilities of even the largest supercomputers. Heuristic methods are therefore commonly used. For over a decade, the Rosetta software suite has been one of the leading software packages for protein design and structure prediction (*1*), using simulated annealing-based heuristics to effect an efficient approach to protein design tasks. Rosetta has been used to design new protein topologies (*2, 3*), large macromolecular assemblies (*4–7*), proteins with the ability to sequester toxic small molecules (*8, 9*) or to bind to other proteins of therapeutic interest (*10, 11*), and enzymes able to catalyze reactions that no known natural enzyme can catalyze (*12, 13*). More recently, Rosetta has been generalized to allow the design of diverse synthetic heteropolymers that fold as proteins do, but which are built from moieties not used by natural proteins (non-canonical side-chains and backbone chemistries) (*14–18*). Non-canonical peptides and peptidomimetics have shown wide utility in biological chemistry and drug design applications (*19, 20*).

Rosetta’s primary design heuristic, called the *Packer*, solves the sequence design problem using simulated annealing-based searches of rotamer space, which, although not guaranteed to converge to the global optimum, tend to find high-quality solutions near the optimum very rapidly (*1, 21*). Unfortunately, the rotamer space quickly grows too large for simulated annealing approaches as the number of designable positions (*N*) or the number of rotamer possibilities at each position (*D*) grows. Large protein design tasks (which have large *N*) or non-canonical design tasks with many choices of chemical building-blocks (which have large *D*) can rapidly become intractable. The *Packer*’s simulated annealing approach is also very sensitive to the shape of the energy landscape, relying on broad energy wells for which a downhill path to the lowest-energy state exists, and sometimes failing to find solutions in narrow energy wells. Alternative approaches, such as dead-end elimination or branch-and-bound searches, have also been used (*22–26*), though these are typically too slow for most design tasks.

Quantum computing provides an attractive alternative. Where classical computers typically solve difficult combinatorial problems by iterating through many possibilities, either exhaustively or using stochastic search heuristics or deterministic optimization algorithms, quantum computers have the potential to represent all possible solutions to a posed problem as a superposition of quantum states (*27, 28*). A quantum algorithm, such as quantum annealing, can then shift the probability distribution of the superimposed states to make the state corresponding to the global optimum overwhelmingly probable on measurement (*29*). The major advantage of a quantum computing approach is the massive parallelism that can be achieved by modelling many solutions simultaneously: indeed, the number of solutions that can be modelled simultaneously doubles with each additional quantum bit, or qubit, added to the system, allowing scaling far beyond anything achievable with classical computers for certain classes of search problems.

Here, we show that the rotamer optimization problem — the central problem that must be solved when designing a protein — maps well to the D-Wave quantum annealer. We demonstrate that this mapping can be made without simplifying the design task or sacrificing the accuracy of the existing classical methods, ensuring that the quantum approach will be at least as useful as current classical approaches. Using classical folding simulations, we also show that the output from our quantum design algorithm, which we call the *QPacker*, has scientific validity comparable to the output from Rosetta’s *Packer*. Because the *QPacker* is a direct mapping of the Rosetta *Packer* to the quantum architecture, it can be applied to any design task to which Rosetta can currently be applied. Since quantum algorithms possess inherently better scaling than their classical counterparts, as larger quantum computers are introduced, we anticipate that the *QPacker* will allow us to tackle larger design tasks than will ever be possible on classical hardware.

## Materials and Methods

### *QPacker* algorithm

To develop the *QPacker*, we first looked to the classical approaches for design tasks. Rosetta’s *Packer* uses simulated annealing-based searches of rotamer space, in which moves involve substituting one rotamer at a randomly-chosen designable position for another allowed rotamer at that position. Moves are rejected or accepted based on the Metropolis criterion. The objective function optimized during this process is typically an approximation of the conformational energy, with a functional form that is rotamer-level pairwise-decomposable. Given *N* designable positions with rotamers at the *N*^th^ position indexed as *r*_*N*, 1_ through 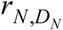, a particular solution is given by a vector 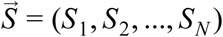, where *S*_1_ through *S*_*N*_ are the indices of the chosen rotamers at each position. The energy of the solution, 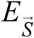, is given by:

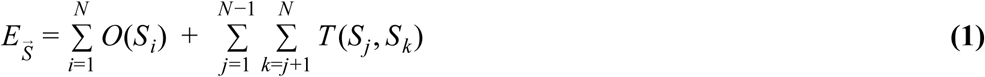

In **Equation 1**, above, *O*(*S*_*i*_) represents a one-body energy of rotamer *S*_*i*_, and *T*(*S*_*j*_, *S*_*k*_) represents a two-body interaction energy between rotamers *S*_*j*_ and *S*_*k*_. Although the functional forms of *O*(*S*_*i*_) and *T*(*S*_*j*_, *S*_*k*_) are very complicated (see (*30*) for a comprehensive review of the Rosetta energy function), the requirement of rotamer-level pairwise-decomposability ensures that the energy calculation can be carried out in a pre-computation, scaling at worst as *O*((*ND*)^2^) (and more commonly as *O*(*ND*^2^), since in practice geometric constraints ensure that the number of interacting pairs of positions tends to scale roughly linearly with the number of designable positions). During the subsequent annealing phase, energies can be updated rapidly when rotamer substitution moves are made by looking up precomputed one- and two-body interaction energies for the affected rotamers only.

The separability of the problem means that the classical simulated annealing phase can be replaced with a quantum annealing phase. Quantum annealing is a hardware-realised metaheuristic approach that minimizes a given objective function over a set of candidate solutions, using the principle that adiabatically changing a quantum-mechanical system that is in its ground state does not perturb the system from its ground state. Finding the global minimum of a posed optimization problem can be formulated as adiabatically evolving the ground state of an initial Hamiltonian, *H*_*S*_ (with a known and easy-to-prepare ground state), to the ground state of the target Hamiltonian, *H*_*T*_ (for which the quantum-mechanical ground state corresponds to the minimum of the classical objective function being optimized). Observational samples can be drawn from *H*_*T*_ to find the minimum of the classical objective function. Assuming that the lowest-energy classical state is highly probable, this procedure can find the global minimum with a finite and hopefully small number of samples. Formulated as a transverse Ising model, the transition between states is controlled by slowly changing the transverse field (*29*). The overall Hamiltonian, parameterized with a value τ that varies from 0 at the start of the annealing phase to 1 at the end, is given by:

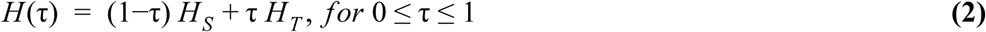

When τ = 0, the lowest-energy state (typically an equal superposition of all states) is easily prepared, and gives all classical configurations equal probability. When τ = 1, the system corresponds to the ground-state Ising problem that one wishes to solve. If the system transitions from τ = 0 to τ = 1 sufficiently slowly, then the solution is guaranteed with high probability; however, the temperature of the system, the influence of external sources of thermal or electrical noise, and the nature of *H*_*T*_ all influence the trade-off between speed and accuracy. Quantum annealing is particularly useful for finding solutions to posed optimization problems in which the search space is discrete, with many local minima, as is the case for rotamer optimization tasks. Current-generation quantum annealers, such as the D-Wave 2000Q system, allow posed problems to be expressed as quadratic unconstrained binary optimization (QUBO) tasks, which involve an objective function *f* with a functional form very similar to that of the Rosetta energy function. Given *n* qubits that, on measurement, can take values of 1 or 0, yielding a state 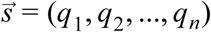 (where *q*_*n*_ represents the value of the *n*th qubit), the functional form of the objective function for a measured state 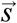 is:

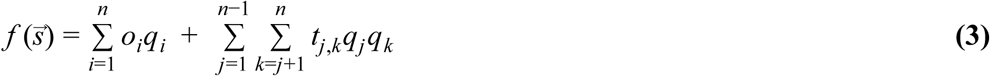

In the above, *o*_*i*_ represents the one-qubit penalty for *q*_*i*_ being 1, and *t*_*j,k*_ represents the two-qubit penalty if *q*_*j*_ and *q*_*k*_ are both 1. The D-Wave system is programmed by setting the values of the *o*_*i*_ and *t*_*j,k*_ coefficients. The quantum annealing algorithm then seeks the state 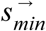 which minimizes the function 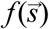.

Note the similarity between **Equations 1** and **3**. Given this, the *QPacker* algorithm can be developed simply by assigning each rotamer under consideration to a different qubit, and then applying three simple rules. First, each *o*_*i*_ must be set to be the classically pre-computed one-body Rosetta energy for that rotamer. Second, each *t*_*j,k*_ for qubits *j* and *k* representing rotamers at *different* sequence positions must be set to be the classically pre-computed two-body Rosetta energy for that pair of rotamers. And third, each *t*_*j,k*_ for qubits *j* and *k* representing different rotamers at the *same* position must be assigned a large positive value (effectively prohibiting solutions in which more than one rotamer is selected at a given position). When the D-Wave quantum processing unit (QPU) is programmed in this way, a Rosetta design task is translated without distortion or simplification into a quantum annealing problem.

### Embeddings on the D-Wave 2000Q

Solving a posed optimization problem on a quantum annealer requires first mapping the optimization task as a graph onto the QPU; however, qubits on current quantum annealers are not fully connected, but rather arranged in particular architectures with each qubit connected to a limited number of adjacent qubits. The D-Wave 2000Q’s “Chimera” architecture is composed of sets of connected unit cells, each with two sets of four qubits in which all of the qubits in the first set is connected to all of the qubits in the second *via* couplers (bipartite connectivity). Additionally, each qubit is connected to two additional qubits in neighbouring unit cells. Unit cells are tiled vertically and horizontally with these inter-unit connections, creating a lattice of sparsely connected qubits. Because of this particular architecture, most optimization problems require minor embedding. For example, the D-Wave 2000Q system does not natively support *K*3 graphs, but these can still be embedded using a penalty model. Because the D-Wave 2000Q system has fewer physical connections than there are pairs of qubits (each qubit being connected to six neighbours), rotamers with more than six interactions must ultimately be represented by more than one qubit, with strong coupling constraints between pairs of qubits representing the same rotamer to ensure that the state is the same when measured. We used the *minorminer* algorithm, which heuristically seeks a minor embedding of a graph representing a dense QUBO in a sparse target graph.

### Decomposing large design tasks

Due to the limited number of qubits and sparse connectivity available on D-Wave’s 2000Q system, the largest graph optimization task that can be embedded onto the system is approximately *K64*. Solving larger design tasks requires using decomposition techniques that effectively break large tasks into smaller segments that can run individually on the QPU, then combine these solutions into an overall solution to the larger task. We used D-Wave’s *qbsolv* algorithm to solve large design tasks, restricting the solver limit to 50. *qbsolv* is a hybrid classical/quantum algorithm that uses a two-level approach. The first level considers the full QUBO instance, and the second level considers a sub-QUBO sized to fit on the underlying D-Wave QPU. The algorithm iterates through trials, with each trial consisting of a set of calls to a sub-QUBO solver for global optimization of the sub-QUBO, and a call to a tabu search algorithm for local minimization (*31–34*). This approach allows the current quantum hardware to effectively solve large combinatorial optimization problems despite the limited number of qubits and connections.

### Modifications to Rosetta

We added a module to Rosetta called the *InteractionGraphSummaryMetric* to allow precomputed *Packer* interaction graphs to be converted to an ASCII format easy for D-Wave scripts to read, and easy to store on disk. The output format contains a full description of the backbone to be designed, the identity and conformation of each rotamer allowed at each position, and the precomputed one- and two-body interaction energies for each rotamer and pair of rotamers, respectively. To permit solutions from the D-Wave system to be imported and converted into an all-atom representation of the designed molecule, we also added a Rosetta module called the *ExternalPackerResultLoader*. This accepts a description of a design task (previously written with the *InteractionGraphSummaryMetric*) and a vector of rotamer selections (one rotamer index per designable position), from which it generates a structure of the design with the selected rotamers placed.

Past studies comparing the Rosetta packer to other methods have been hindered by the opacity of the Rosetta source code, and have struggled to ensure that identical design tasks were solved by Rosetta and other methods. Separating the performance of the interaction graph precomputation from the simulated annealing runs was also a challenge. In order to carry out direct comparisons of the Rosetta *Packer*, the *QPacker*, and *Toulbar2*, we added a Rosetta application called *compare_external_packer_output*. This application accepts as input a design task definition (as written by the *ExternalPackerResultLoader*), a solution vector (computed by an external algorithm such as the *QPacker*), and a value for the number of Rosetta replicates to perform. It provides the design task from the definition file to the Rosetta *Packer*’s simulated annealer, timing each simulated annealing replicate and writing the resulting structures, energies, and runtimes. For convenience, it also converts the external packer solution to a full structure. Finally, it writes the same design task again, this time as a *Toulbar2* `.cfn` file, allowing it to be solved with that application as well.

All of these modules and applications are currently available in Rosetta git revision `cae61325ed5d03f361826d4a5c3160a87c708f27`, and will be incorporated into the master branch of Rosetta in the near future. In the meantime, the source code is available for academics, nonprofit users, and governments on request.

### Test applications

Peptide and protein design tasks range from small computational tasks with only dozens of solutions, which can be solved by exhaustive enumeration, to massive computational tasks with astronomical numbers of solutions, which can exceed the capabilities of stochastic methods like simulated annealing. In order to test the *QPacker*, we devised three categories of design task. First, as a scalable test case, we used Rosetta’s parametric design tools (*18, 35, 36*) to construct helical bundles of various sizes, then set up design tasks involving core positions, and using different sets of rotamers. We allowed increasing numbers of core positions to pack using all rotamers of one of the 18 rotameric canonical amino acids for each design task (excluding alanine and glycine, which have no rotamers). We generated 400 design tasks with the number of designable positions, *N*, ranging from 1 to 40, and the geometric average number of rotamers per position, *D*, ranging from 1.03 to 6.77. These tasks were deliberately designed to fit on the D-Wave QPU. Each design task was carried out using the *QPacker*, as well as 100 times using the Rosetta *Packer*, to measure performance (computation time). We also solved each design task using the exact branch-and-bound solver *Toulbar2* (*25, 26*), *and compared Packer* and *QPacker* solution convergence to the global optimum. To provide a reference for anticipated future *QPacker* performance, we also solved 5,376 additional design tasks that were too large for a single D-Wave QPU (with *N* up to 40 and D up to 54.0) using the Rosetta *Packer* and *Toulbar2*.

Second, to demonstrate the *QPacker’s* utility, we applied the algorithm to realistic peptide macrocycle design tasks. As a test case, we chose an exotic peptide secondary structure almost never seen in nature, the α-sheet. The α-sheet is only stable when built from alternating mixtures of D- and L-amino acid residues (*37, 38*), and it represents an interesting design challenge for any design algorithm. We classically sampled 16-residue peptide macrocycle α-sheet backbone conformations using a protocol developed in the RosettaScripts scripting language (*39*), then carried out design on the quantum annealer.

Finally, we sought to apply the *QPacker* to more protein-like design tasks, in a size range in which a hydrophobic core is possible. Although the current quantum annealing hardware limits the size of design tasks that can be tackled, molecules with internal *n*-fold symmetry can have *n*-fold more amino acid positions without increasing the number of *unique* designable positions or the overall computational complexity of the design task (*40, 41*). We therefore sampled 32-residue S2-symmetric macrocycle conformations classically, and designed S2-symmetric molecules with hydrophobic cores and polar surfaces using the *QPacker*. Here again, we deliberately chose an exotic design challenge: natural proteins, built from amino acids with the same chirality, cannot access improper rotational symmetries involving mirror operations. The topologies on which we focussed consisted of helices of opposite handedness packing against one another, representing a tertiary motif found in no natural protein. Despite the exotic topology, the principles of protein folding — namely, that well-folded molecules possess well-packed, hydrophobic cores — still apply.

## Results

### Performance scaling

We carried out performance benchmarks on 400 helical bundle design tasks that could fit on the D-Wave 2000Q QPU. Each task was solved on the D-Wave 2000Q system, as well as with the classical Rosetta *Packer* and with the branch-and-bound solver *Toulbar2*. We also applied the two classical methods to an additional 5,376 design tasks that were larger than what could fit on the D-Wave QPU. In each case, the total time to solution was measured. On the quantum annealer, this all-inclusive runtime includes the time to load the design task onto the quantum QPU, and the annealing time, read time, and reset time multiplied by the number of replicates needed (due to electronic and thermal noise, which disrupts a subset of runs). For the classical methods, this included the time needed to set up the design task in memory in a form that could be solved, the time to carry out the simulated annealing (for the Rosetta *Packer*) or the branch-and-bound search (for *Toulbar2*), and the time to convert the solution in memory to a form suitable for output. Precomputation of the interaction graph was excluded from the calculation, since this was common to all three methods; likewise, disk reads, network transfer times, and disk writes, which varied across the methods but are not intrinsic to any of the methods, were excluded. Empirically, we found that both classical approaches scale with a power of the complexity of the design task, as expressed as the product of the geometric average of the number of rotamers per position and the number of positions. Notably, for problems in this size range, the *QPacker* achieves nearly constant performance, independent of the complexity of the task, up to the limits of the quantum computer’s size (**Fig. 1**). The limited size of the 2,048-qubit QPUs that are the state of the art currently prevents the *QPacker* from outperforming the classical approaches. Larger design tasks can be decomposed using the *qbsolv* hybrid algorithm, but beyond the size of the QPU, runtimes cease to be constant and classical scaling applies. Note that, at the time of writing this manuscript, the next-generation D-Wave QPU, named “Pegasus”, has been announced. According to the manufacturer, this QPU will feature 5,640 qubits, and will raise the number of connections per qubit from 6 to 15 (*42*), which would raise the limit on the size of design task to which the *QPacker* could be applied. It remains to be seen exactly how the *QPacker* scales on larger quantum computers; however, we anticipate better than classical scaling.

**Figure 1:**
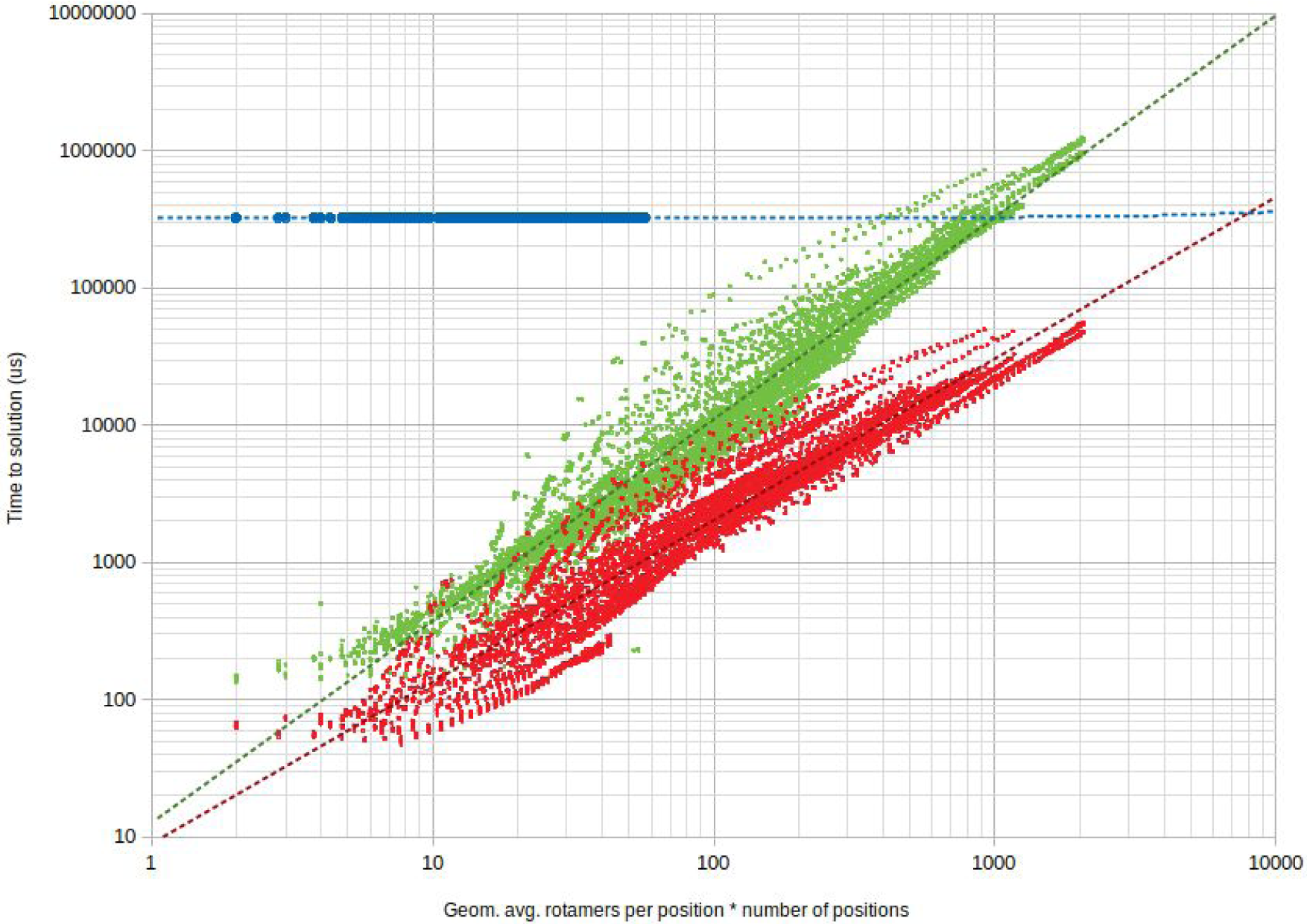
Comparison of performance of the quantum design algorithm (*QPacker*, blue points) with a classical, exact branch-and-bound algorithm (*Toulbar2*, green points) and the classical simulated annealing method used by the Rosetta *Packer* (red points). 400 helical bundle design tasks that fit onto the D-Wave 2000Q QPU (solved using the *QPacker*, the Rosetta *Packer*, and *Toulbar2*), and another 5,376 design tasks that did not (solved using the Rosetta *Packer* and *Toulbar2*), were used for this analysis. The vertical axis shows time to solution in microseconds, with lower values indicating better performance. A proxy for the complexity of the search space, the product of the geometric average of the number of rotamers per position and the number of designable positions (which is approximately equal to the total number of rotamers), is shown on the horizontal axis. The quantum annealing-based approach takes approximately constant time over the range of problem sizes that could be tested, while both the branch-and-bound method and the simulated annealing method scale exponentially. Dashed lines show a linear fit to the *QPacker* data and power fits to the *Toulbar2* and Rosetta *Packer* data.

### Design accuracy and scientific utility

We next explored whether the *QPacker* achieves accuracy comparable to *Toulbar2* or Rosetta’s *Packer*. Examining the pool of 620 32-residue S2-symmetric design tasks, and comparing the pools of designs produced by the classical *Packer* and by the *QPacker* to the pool produced by the exact *Toulbar2* solver, we found that the *Packer* more consistently converged to the global optimum (78.8% of the time) in this size-range than did the *qbsolv*-extended *QPacker* (1.1% of the time). Nevertheless, both methods consistently found solutions close to the global optimum, with the *QPacker*’s solutions falling within 1.16 kcal/mol of the global optimum 10% of the time, 4.35 kcal/mol of the global optimum 50% of the time, and 8.45 kcal/mol 90% of the time. In 2.6% of cases, the *QPacker* found a lower-energy solution than did the Rosetta *Packer*, and both methods found the same solution in an additional 1% of cases. **Fig. 2** shows the close correlations between the energies of the solutions found by the *Packer*, the *QPacker*, and *Toulbar2*.

**Figure 2:**
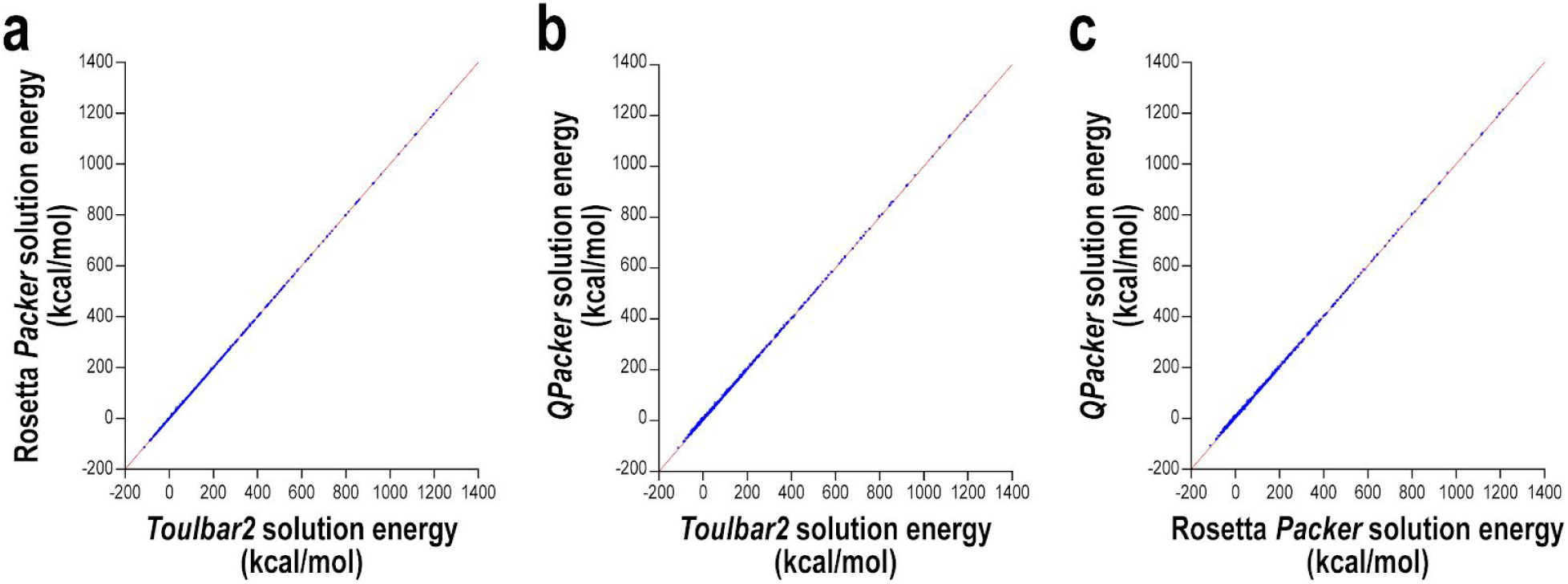
Comparison of energies of solutions found by Rosetta’s *Packer* and the *qbsolv*-based *QPacker* to the global optimum found by *Toulbar2*. Energies for 622 32-residue S2-symmetric macrocycle design tasks are plotted as blue points. The line y=x is shown in red. (**a**) Comparison of energies for 32-residue S2-symmetric designs, produced by the Rosetta *Packer* vs. those of designs produced by *Toulbar2*. Rosetta does not consistently converge to the global optimum in this size-range, as indicated by some points above the line. (**b**) Comparison of energies of designs produced by the *QPacker* vs. those of designs produced by *Toulbar2*. The *QPacker* also does not consistently converge to the global optimum in this size-range, but energies are consistently close to the global optimum. (**c**) Comparison of energies of designs produced by the *QPacker* vs. those of designs produced by the Rosetta *Packer*. Although the *QPacker* tends to find slightly higher-energy solutions than the Rosetta *Packer*, it outperforms the Rosetta *Packer* 2.6% of the time, and matches the Rosetta solution an additional 1.0% of the time.

Convergence to the global optimum does not necessarily correlate with maximization of the energy gap between the designed backbone conformation and all alternative conformations. To assess energy gaps and folding propensity of designs, we carried out classical conformational sampling on each of the 620 designs produced with the *QPacker*, using the Mira Blue Gene/Q supercomputer, and sampling an average of 86,000 conformations for each designed sequence to determine whether the designed conformation was a unique low-energy state, or whether there were alternative low-energy states. For each design, we computed the metric *P*_*near*_ (*16, 17*), which quantifies fold propensity from poor (0) to excellent (1). Given *M* samples, each with a computed energy (*E*_*i*_) and RMSD from the target conformation (*R*_*i*_), *P*_*near*_ can be computed as:

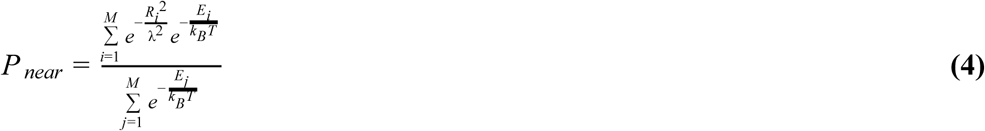

In the above, *λ* is a parameter that determines how closely a conformation must match the designed conformation to be considered “native-like”, which we set to 2 Å. The value of the Boltzmann temperature, *k*_*B*_*T*, was set to 0.62 kcal/mol to approximate the room-temperature thermodynamic distribution of states.

*P*_*near*_ values over 0.9 correlate very strongly with experimental success (*17*). We found that the pool of 622 *QPacker*-produced designs contained 6 high-quality designs with *P*_*near*_ values greater than 0.9, and an additional 15 designs with *P*_*near*_ values between 0.8 and 0.9. A representative classically-computed energy landscape for one of the *QPacker*-produced designs with *P*_*near*_ > 0.9 and a histogram of the fraction of designs in each *P*_*near*_ bin are shown in **Fig. 3**. The observed design success rate (1% over *P*_*near*_=0.9, and 3.4% over *P*_*near*_=0.8) is comparable to that from many *Packer*-based design projects, indicating that the *QPacker* can be used to produce scientifically useful results, and can produce sequences for subsequent wet-lab experiments. Chemical synthesis and experimental characterization of these designs is ongoing work.

**Figure 3:**
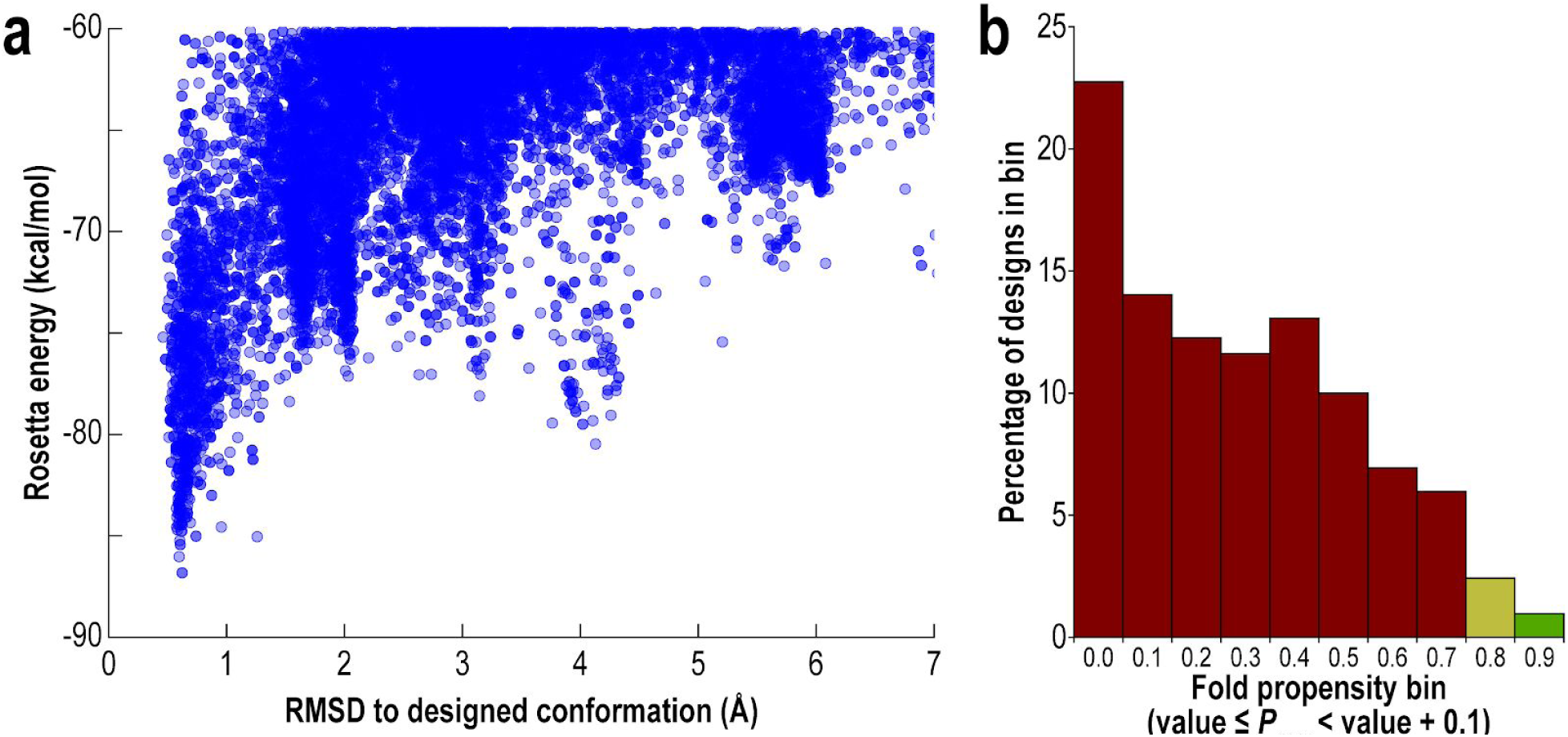
Results of folding simulations performed to provide classical computational validation of peptide designs produced by the *QPacker*. (**a**) A classically-computed energy landscape, produced by large-scale sampling on the Mira Blue Gene/Q supercomputer, for one of the top 32-residue S2-symmetric macrocycle designs produced by the *QPacker*. Each blue point is the result of a single conformational sampling trajectory, plotted as a function of deviation from the designed conformation (horizontal axis) and computed energy (vertical axis). As shown, this design has a unique low-energy state very close to the designed conformation, and all alternative conformations are much higher in energy. (**b**) Histogram showing the design pool binned by energy funnel quality (based on the *P*_*near*_ metric, computed from large-scale conformational sampling runs on Mira). *P*_*near*_ values over 0.9 (green bar) correlate very strongly with success in wet-lab experiments, and *P*_*near*_ values between 0.8 and 0.9 (yellow bar) are often acceptable. As shown, the design pool contains a number of designed molecules that are predicted to fold, with a success rate (fraction of total design pool that is predicted to fold) comparable to that achieved in many classical design projects.

### Qualities of designs produced

Even when peptide and protein design algorithms do not converge to the global optimum, they must still produce designs with favourable structural features amenable to folding. **Fig. 4** shows representative designs from the pools of 16-residue α-sheet designs and 32-residue S2-symmetric coiled-coil designs. As shown, the *QPacker* successfully finds closely-packed arrangements of amino acid side-chains, particularly in the hydrophobic core (orange) of the 32-residue designs. Other favourable features, such as salt bridges between oppositely-charged amino acid residue types, are also in evidence.

**Figure 4:**
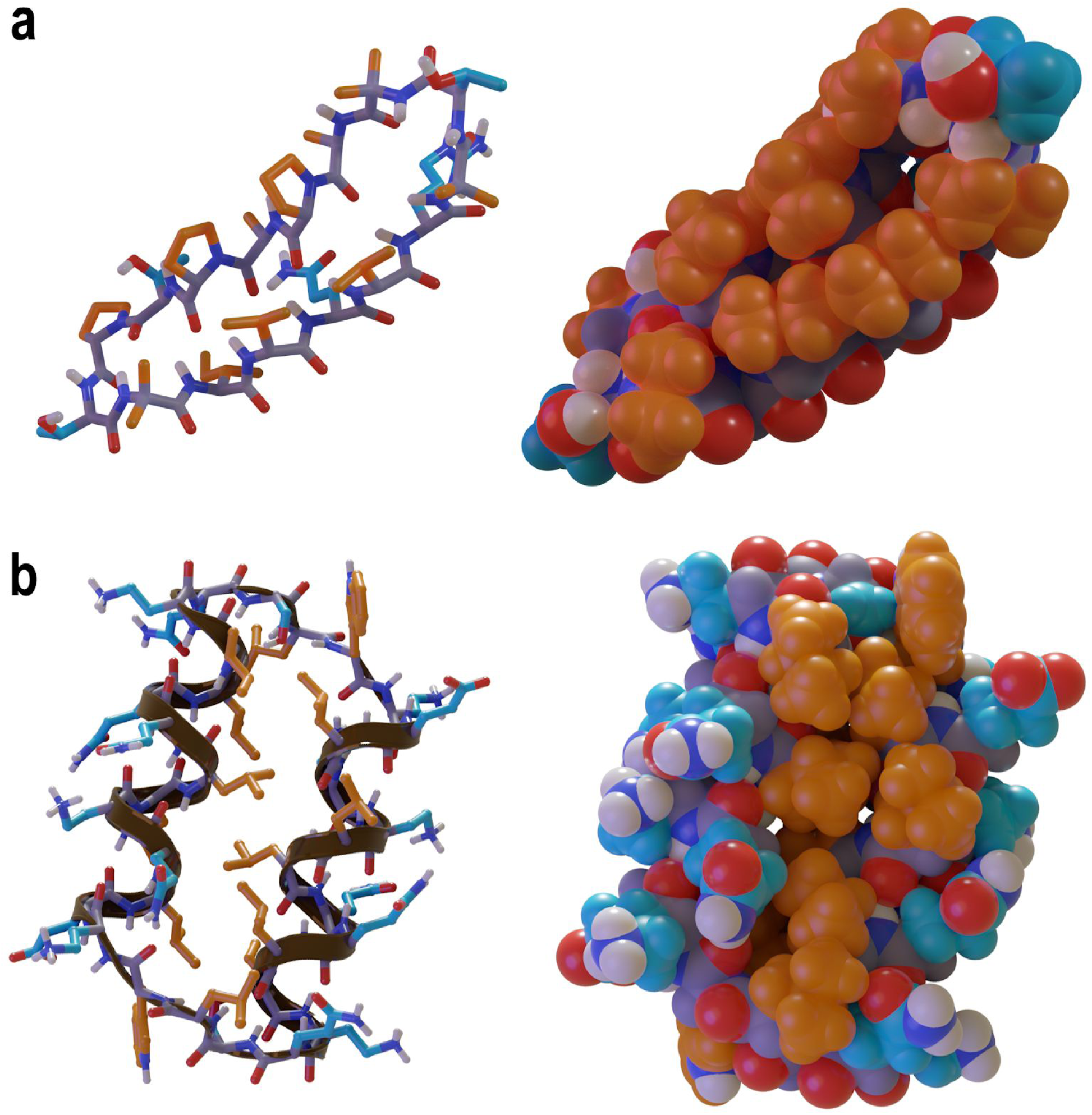
Representative designs produced by the *QPacker*. (**a**) Sticks (left) and space-filling (right) models of a representative 16-residue α-sheet design. Apolar side-chains are shown in orange, polar side-chains, in cyan, and backbone atoms in grey. Nitrogen, oxygen, and polar hydrogen atoms are shown in blue, red, and white, respectively. The *QPacker* consistently found solutions with good side-chain packing, particularly between apolar groups. (**b**) Ribbon and sticks (left) and space-filling (right) models of a representative 32-residue S2-symmetric coiled-coil design. Colours are as in the previous panel. The excellent packing of the hydrophobic core is evident. Other features important for folding, such as salt bridges and side-chain hydrogen-bonding interactions, are also in evidence.

## Discussion

For a protein designer choosing whether to use quantum annealing as a design tool, there are two obvious questions. First, when can we anticipate that there will be an advantage to using the *QPacker* over existing classical methods? And second, can the *QPacker* produce results of sufficient accuracy to be useful for informing design projects and guiding wet-lab experiments?

In considering the first question, it is important to differentiate quantum *supremacy* (the demonstration that a quantum computer can solve a problem, useful or not, that a classical computer cannot (*43, 44*)) from quantum *advantage*. Showing quantum supremacy is beyond the scope of this work, and is properly in the domain of those working to advance the hardware; the reader is directed to recent high-profile publications on the subject for more information (*45–48*). Quantum advantage, on the other hand, is commonly used to mean the demonstration that a problem may be solved more quickly using a quantum computer than a classical computer, or, more broadly, that there exists a practical reason for choosing to solve a useful, real-world problem on a quantum computer rather than on a classical computer. By the latter definition, this can represent a speed advantage, an advantage in the properties of solutions or distribution of samples produced, or other advantages such as energy consumption (*49–51*).

In the area of protein design, quantum advantage could be realized by either improvements to the hardware (*e.g.* having more qubits, lower noise, or increased connectivity), or by improvements to the algorithms that run on these devices (*e.g.* denser and more efficient mapping to the quantum device). There are many unknowns involved in trying to predict when there would be a quantum advantage for protein design, not the least of which is the question of whether such favourable scaling as we have observed will continue on future-generation systems. But considering only improvements to the hardware, and supposing that the observed scaling of the current algorithm continues, by naïve extrapolation of the lines in **Fig. 1** to the points of intersection it is possible to estimate that given a quantum annealer of sufficient size to handle about twentyfold more rotamers, the *QPacker* would compete with the exact branch-and-bound approaches, and given a quantum annealer able to handle a hundredfold more rotamers, the *QPacker* would compete with Rosetta’s *Packer*. With complete connectivity, this would require machines with twentyfold or a hundredfold more qubits, respectively. However, the limited connectivity of the D-Wave system’s qubits allow it to simulate a set of fully-connected qubits with the number of simulated, fully-connected qubits scaling approximately with the square root of the number of physical qubits. If we were to assuming pessimistically that there will be no improvements made in connectivity (which would allow larger design tasks to be handled with fewer qubits), and that no reductions in thermal or electronic noise will be achieved (which would allow design tasks to be handled faster, with less repeat sampling, shifting the blue points in **Fig. 1** downward and allowing them to intersect the red and green lines sooner), quantum advantage over *Toulbar2* and the Rosetta *Packer* would be expected with machines that are approximately 400 and 10,000 times larger than the current-generation machines, respectively. Currently, we are in a Moore’s law period of increase in the size of quantum computers: for the past decade, the number of qubits in adiabatic quantum annealers like the D-Wave system has doubled approximately every year, which alone would result in a 400-fold size increase in about 8.6 years, and a 10,000-fold size increase in about 13 years. However, major strides are also being made in improving the connectivity and noise of these systems. These advancements can reasonably be anticipated to allow quantum advantage for protein design tasks in a considerably shorter period. Indeed, a next-generation D-Wave QPU has already been announced which increases the degree of qubit connectivity from 6 to 15, and claims have been made of noise improvements (*42*). Since current annealing times are on the order of 20 µs, in the very best case, improved noise levels eliminating the need for repeated sampling would bring the *QPacker*’s speed on the problems that it can handle now to under that of the Rosetta *Packer* or *Toulbar2* without any increases in size or connectivity. Given plausible increases in connectivity and reductions in noise, quantum advantage could be realized in the near term (2-5 years). Further improvements to the mapping algorithm or development of new classical-quantum hybrid approaches are also likely to shorten the time.

We have defined quantum advantage to mean that the quantum computer must be able to solve a *useful, real-world* problem, and do so well enough to make it desirable to use the quantum computer instead of a classical computer to solve that problem. This means that the quantum computer must produce useful, accurate results, which brings us to the second question: can the *QPacker* produce results of sufficient accuracy to inform wet-lab experiments? In considering this question, “accuracy” can have two meanings. First, an accurate design method should find solutions at or near the global optimum. *Toulbar2*’s branch-and-bound approach is guaranteed to find the global optimum (*25*), while the Rosetta *Packer*’s stochastic approach is not (*1, 21*), but nonetheless tends to find solutions close to the global optimum in much less time. In this sense, the accuracy of the *QPacker* is comparable to that of the *Packer*, though the *Packer* currently converges to the global optimum more frequently. Primary reasons for the *QPacker* failing to converge to the global optimum are noise (individual quantum annealing runs fail to find the lowest-energy state) and the *qbsolv* decomposition (the outer, classical sampling is not guaranteed to converge). The anticipated hardware improvements discussed above — reduction of noise, and increases in size and connectivity that will reduce reliance on *qbsolv* — are both expected to improve this accuracy.

“Accuracy” can also refer to the scientific utility of the result — particularly, the ability of the method to produce designed sequences that are predicted to fold into the designed conformation. Folding requires the *unique* stabilization of the designed conformation, and the destabilization of all alternative conformations, maximizing the energy gap between folded and all alternative states (*43–45*). Because the formulation of a design task as a packing problem considers only the polypeptide backbone conformation that one seeks to stabilize, and disregards the ensemble of alternative conformations that the molecule might access (regardless the optimization method employed), designers necessarily take a “guess-and-check” approach, first creating a large pool of designs by relatively computationally inexpensive heuristic methods (like the Rosetta *Packer* or the *QPacker*), then checking each by relatively computationally expensive structure prediction algorithms. The latter class of algorithms, which include Rosetta’s *AbinitoRelax* (*46–49*) and *simple_cycpep_predict* applications (*16, 17*), sample tens of thousands to millions of alternative backbone conformations given a fixed sequence, relaxing and computing energies of each conformation sampled. The ensemble is then analyzed to determine whether the sequence *uniquely* favours the designed conformation (in which case the design is typically selected for wet-lab experiments), or whether it has alternative low-energy conformations (in which case the design is rejected), with metrics like *P*_*near*_ serving as useful means of quantifying the quality of the energy funnel. Because there is no guarantee that the unique optimal solution to the packing problem is the sequence that creates the largest energy *gap* between the designed conformation and all alternative conformations, many solutions near the optimum are of interest; that is, a method that produces near-optimal solutions to the packing problem, but which rarely converges to the global optimum, can still be useful for finding sequences that fold. In this sense, we have demonstrated that the *QPacker* has scientific utility, in that design pools comparable in size to those produced classically (on the order of hundreds or thousands of designs) contain sequences which, when analysed with expensive, rigorous validation algorithms, are predicted to fold. The observed success rate from computational validation, on the order of 1%, is comparable to that in many classical protein design projects.

We have shown here that one of the best-performing classical approaches to protein design, the Rosetta software suite’s *Packer*, can be mapped to existing quantum computers. Moreover, we have demonstrated that the mapping can be performed without simplifying the problem or abandoning heavily optimized aspects of the Rosetta package, such as its high-accuracy energy function and its backbone-dependent rotamer libraries. Although the noise levels and size of current quantum hardware is a limitation, hybrid algorithms such as *qbsolv* extend the utility of the approach to problems of considerable complexity. Moreover, the method can produce scientifically useful results, such as the generation of pools of sequences which contain high-interest sequences predicted to fold into the desired structures, informing subsequent wet-lab experiments. As quantum hardware grows and noise levels fall, we anticipate that the *QPacker*’s remarkable scaling will permit design tasks that are currently intractable on classical hardware.

## Acknowledgements

This work was supported by the Simons Foundation (V.K.M., D.R., and R.B.). A.M.W. is supported by an ARO MURI sub-award. H.I.M. and P.S.A. are supported by NIH grant R35GM130333. H.I.M. is supported by the Kramer Predoctoral Fellowship. S.S. is funded by NASA ROSES AIST NNH16ZDA001N-AIST. An award of computer time was provided to V.K.M. by the Innovative and Novel Computational Impact on Theory and Experiment (INCITE) program. This research used resources of the Argonne Leadership Computing Facility, which is a DOE Office of Science User Facility supported under Contract DE-AC02-06CH11357. The authors would also like to thank D-Wave Systems, Inc. and the Creative Destruction Lab at the University of Toronto for useful discussions and support.

## Footnotes

1 In design tasks with different numbers of allowed side-chain possibilities at each position, this relationship still holds, with D representing the geometric average number of possibilities per position.

